# Niclosamide Prodrug Enhances Oral Bioavailability and Targets Vasorin-TGFβ Signaling in Hepatocellular Carcinoma

**DOI:** 10.1101/2024.10.15.618538

**Authors:** Mingdian Tan, Wei Ye, Yi Liu, Xiaowu Chen, Lakshmi Huttad, Mei-Sze Chua, Samuel So

## Abstract

**Background:** Hepatocellular carcinoma (HCC) ranks third in cancer-related deaths worldwide, with limited therapeutic options. While niclosamide (NIC) has shown potential for repurposing in HCC, its poor water solubility and low bioavailability limit its efficacy, and its mechanisms of action are not yet fully elucidated.

**Methods:** We designed a water-soluble NIC prodrug (NIC-PS) and evaluated its efficacy through **in vitro** and **in vivo** studies, including pharmacokinetic (PK) and pharmacodynamic (PD) assays, HCC patient-derived xenograft (PDX) models were applied in two independent experiments, vasorin (VASN) knockout models, and combination treatments with NIC-PS and sorafenib or anti-PD-L1 antibody. Bioinformatic analyses and western blotting were used to investigate NIC-PS’s target, VASN, and related signaling pathways.

**Results:** NIC-PS exhibited a **ten-fold increase in oral bioavailability** and reduced tumor volume by over **75%** in HCC PDX models. NIC-PS directly binds and suppresses VASN, suppressing TGFβ signaling and reducing SMAD2/3 phosphorylation. VASN inhibition led to a **50% tumor reduction**, and NIC-PS enhanced responses to sorafenib and anti-PD-L1 therapy.

**Conclusions:** NIC-PS, equal to 36% of NIC in molecular weight, offers improved bioavailability, efficacy, and a **novel mechanism of action in targeting VASN**, showing promise for HCC treatment alone or in combination therapy.

## Background

Hepatocellular carcinoma (HCC) is the sixth most prevalent cancer globally, and is the third leading cause of cancer-related deaths^1^. Partly due to the lack of sensitive early detection methods, HCC patients often present with symptoms when the disease is advanced, thereby limiting treatment options^2^. Additionally, treatment intervention must consider complex and inter-related pathologies underlying HCC, such as liver inflammation, viral infections, cirrhosis, and compromised liver function. Additionally, HCC has vastly heterogeneous molecular features, which poses significant challenges in developing effective therapeutic strategies for HCC^3^. Despite the availability of multiple therapeutic options ^4^, none have demonstrated significant efficacy in alleviating the disease burden on patients, or enhancing their quality of life. Consequently, the mortality and recurrence rates of HCC remain high, and there exists an unmet need in developing novel complementary approaches to effectively manage this typically fatal malignancy.

Our lab previously undertook a bioinformatics approach to repurpose drug candidates for treating HCC, which yielded niclosamide (NIC), an anti-parasitic drug that is approved by the Food and Drug Administration for treating tapeworm infection^5^. For this indication, NIC acts as an uncoupler of oxidative phosphorylation in the mitochondria to impede metabolism^6^. Cumulative research revealed that NIC affects multiple signaling pathways and biological processes critical in multiple solid and non-solid cancers, and other diseases (such as bacterial and viral infection including Covid-19, metabolic diseases such as Type II diabetes, NASH and NAFLD, artery constriction, endometriosis, neuropathic pain, rheumatoid arthritis, sclerodermatous graft-versus-host disease, and systemic sclerosis)^7^. In cancer, NIC and its analogs have been shown to inhibit cancer cell growth through multifaceted pathways, such as the disruption of signaling cascades (including Wnt/β-catenin, mTOR, and STAT3 pathways^5^, NF-κB pathways^8^), and affecting cellular processes (such as apoptosis^9^, autophagy^10^, cell cycle arrest^11^, disrupting mitochondrial function and increasing reactive oxygen species^12^, modulating the tumor microenvironment^13^ and immune modulation^14^). Additionally, NIC exerts important immunomodulatory actions on specific subsets of immune cells and related pathways^15^.

Despite encouraging *in vitro* data and mechanistic rationale of its anti-tumor potential, recent clinical trials of NIC in colon and prostate cancers have not been unsuccessful (NCT02687009; NCT03123978; NCT02519582; NCT02532114). This is attributed to poor systemic exposure due to low water solubility and consequently low bioavailability, preventing NIC from reaching the solid tumor sites. Several approaches have been evaluated to overcome these limitations; for example, the formation of cocrystals^16^, reformulations based on polymeric micelles^17^ and polypeptide nanoparticles^18^, self-microemulsifying drug delivery system encapsulating NIC^19^, and the design of derivatives with enhanced water solubility, including NIC ethanolamine salt (NEN)^20^, novel NIC analogs^21^, and phosphate^22^ and valine prodrugs of NIC^23^. In particular, the prodrug approach has proven to increase systemic exposure to NIC to achieve desired therapeutic effects with minimal toxicity in the disease models studied^22,23^.

In this study, we designed a water-soluble prodrug derivative, termed NIC-PS, in an attempt to facilitate the clinical translation of NIC for use as a therapeutic option in HCC. We evaluated its anti-tumor activities in HCC cell lines and an orthotopic patient-derived xenograft (PDX) mouse model of HCC, as well as biological mechanism of action.

## Methods

### Culture of HCC cell lines

Human HCC cell line Hep3B, HepG2, SNU398, SNU423, and SNU449 and normal liver cells THLE2, were purchased from American Type Culture Collection (ATCC), whereas Huh7 was a gift from Dr. Mark Kay (Stanford University). Huh7 cells were cultured in Eagle’s Minimum Essential Medium Medium (cat# 30-2002, ATCC); HepG2 and Hep3B cells in Dulbecco’s Modified Eagle’s Medium (cat# 11095098, Gibco); SNU398, SNU423, and SNU449 were cultured in RMPI 1640 medium; THLE cells were cultured in BEGM (CC3170, Lonza Biologics). All media were supplemented with 10% fetal bovine serum (FBS), 100 U/mL penicillin, and 100 μg/mL streptomycin. Cells were maintained at 37 in a humidified 5% CO_2_ atmosphere.

### Cell proliferation assay

The effect of NIC, NEN, and NIC-PS on the growth of HepG2 and Huh7 cells were assessed using the CellTiter 96 Aqueous One Solultion Cell Proliferation assay (cat# G4100, Promega). Cells were seeded at 3×10^3^ cells per well in 96-well plates and allowed to adhere overnight. Cells were then treated with NIC, NEN, and NIC-PS at a concentration range of 0.033 µM to 50 µM for 72 h, then Optical density values at 570 nm were measured using a SYNERGY-LX absorbance microplate reader (BioTek) based on manual protocol. The IC_50_ values were calculated through nonlinear curve fitting using GraphPad Prism 10.0 software. For co-treatment studies, NIC-PS was co-incubated with either anti-PD-L1 (10 μg/ml, Cat# SF63, Sino biological) or sorafenib (5 μM, cat # S7397, Selleckchem) for 72 h prior to measurement of cell viability using the MTS assay. For anti-PD-L1 co-treatment assay, IgG (10 μg/ml, SinoBiological) was used as the control.

### Culture of primary hepatocytes and cell viability assay

Primary human hepatocytes were obtained from Sekisui XenoTech (cat# CHP48), and cultured in 48-well plates using specified culture medium (cat# K8200) according to vendor protocols. Following cellular adherence, cells were treated with a range of concentrations of NIC, NEN, or NIC-PS for 72 h, with a medium replacement every 24 h. Cell viability was assessed after 72 h, using the CellTiter-Glo® 2.0 Cell Viability Assay solution (cat# 9243, Promega) according to the manufacturer’s instructions.

### Pull-down assay and mass spectrometry

To identify potential protein targets that may bind to NIC, we used a pull-down assay as previously described^24^. Briefly, NIC was immobilized to epoxy-activated-Agarose Sepharose 6B resin bead (Cat# E6754, Millipore Sigma), and the resulting NIC resin beads were incubated with HepG2 whole cell lysate for 4 h. The complex of bound protein and NIC resin were precipitated and electrophoresed by SDS-PAGE, using Huh7 cell lysates, NIC-conjugated resin, and blank resin as controls. Protein bands of interest were excised from the gel, and submitted to the Stanford Mass Spectrometry Core Facility for mass spectrometry identification.

### Thermal shift assay

Recombinant human VASN/SLIT-like 2 Fc chimera protein, CF (Cat# 10139-VN-050, R&D Systems) was prepared according to previously described protocol^25^, and diluted to concentrations of 62.5 μg/ml, 125 μg/ml, 250 μg/ml, 500 μg/ml, and 1,000 μg/ml in phosphate-buffered saline (PBS) (consisting of 5 μl VASN protein + 15 μ L PBS + 1 μL 10 μM NEN or 1μl DMSO, made up to a total volume of 50 μL using PBS). Subsequently, it was labeled with the Protein Labeling Kit RED-NHS (cat# MO-L001, NanoTemper Technologies) as per the manufacturer’s protocol, and the assay was done on a Monolith instrument (NanoTemper Technologies).

### Protein extraction, Western blotting

Total protein was extracted from cultured cells using the T-PER Tissue Protein Extraction Reagent (cat# 78510, Thermo Fisher Scientific), and quantified using the BCA Protein Assay Kit (cat# 23225, Pierce). Equal protein amounts (20 μg) were electrophoresed on 4-12% polyacrylamide gels (Invitrogen) and transferred onto polyvinylidene difluoride membranes. The membranes were then blocked using 5% nonfat milk for 1 h and incubated with primary antibodies against VASN (cat# ab156868, abcam, 1:500 dilution), IL33 (cat# A8096, ABclonal, 1:1000 dilution), TGF-β (cat# A15103, ABclonal, 1:2000 dilution), SMAD2/3 (cat# 3102, CST, 1:1000 dilution), p-SMAD2/3 (cat# AP0548, Abclonal, 1:1000 dilution), PD-L1 (cat# 13684S, CST, 1:1000 dilution) or GAPDH (cat# 60004-1-Ig, Proteintech, 1:20,000 dilution). Specific proteins were visualized utilizing a fluorescent secondary antibody, IRDYE 800CW goat anti-rabbit (1:10,000 dilution, cat# 926-32211, LI-COR Biotechnology) or IRDYE 680RD goat anti-mouse rabbit (1:10,000 dilution, cat# 926-68070, LI-COR Biotechnology).

### ELISA assay

Assays for IL33, VASN, and TGFβ were conducted according to the manufacturers’ instructions. Briefly, cell supernatants were collected and concentrated 20-fold using Pierce™ Protein Concentrators PES (cat# 88528, Thermofisher). The concentrated supernatants were then assayed using the IL-33 (cat# TBS3245, Tribioscience), VASN (cat# TBS3246, Tribioscience), and TGFβ (cat# DY240-05, R&D Systems) ELISA kits.

### *In vivo* efficacy study of NIC-PS in orthotopic patient-derived xenografts in mice

All animal studies were performed strictly according to the guidelines and rules concerning laboratory animal care, and were approved by the Institutional Animal Care and Use Committee (IACUC) at Stanford University (Protocol number: APLAC-20167).

Prior to testing the efficacy of NIC-PS in mouse models of HCC, we performed a single dose pharmacokinetic analysis in non-tumor bearing mice, using NIC (40 mg/kg) or NIC-PS (equivalent to 40 mg/kg of NIC). Non-tumor bearing mice (NOD.Cg-Prkdc^scid^ Il2rg^tm1Wjl^ /SzJ (NSG) mice (cat#005557, the Jackson Laboratory; male or female, aged 6 to 10 weeks)) were given a single dose of NIC (40 mg/kg) or NIC-PS (equivalent to 40 mg/kg of NIC) by oral gavage (NIC-PS was dissolved in water, NIC was dissolved in 0.5% Methylcellulose) (n=3 in each group). Plasma was collected at different time points after compound administration (5 min, 15 min, 30 min, 1 h, 2 h, 4 h and 24 h), and processed for analysis of NIC concentration using the liquid chromatography and tandem mass spectrometry (LC-MS/MS) method (Integrated Analytical Solutions, Berkeley, CA).

Orthotopic patient-derived xenografts (PDX) were established as described previously^26^. The PDX-bearing mice were then randomly divided into three groups (n=10 in each group): 1). Control group treated with distilled water; 2). NIC-PS group treated with NIC-PS at 100 mg/kg in water; 3). NEN group treated with NEN at 200 mg/kg NEN in 0.5% methylcellulose.

Treatments were given once daily by oral gavage. Tumor growth was monitored once a week *via* bioluminescence imaging, and body weight was recorded three times a week. Blood samples (50 µL) were collected at 0.5, 4 h after the first dose, and plasma obtained for measurement of NIC concentration using LC-MS/MS (Integrated Analytical Solutions, Berkeley, CA). PDX and adjacent normal liver tissues were harvested at the end of the treatment period, and processed for measurement of NIC concentrations and for immunohistochemical staining. Tumor volumes were calculated using the following equation:

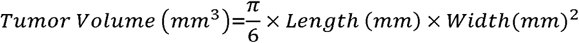

### Immunohistochemistry

Paraffin blocks of harvested tissues were de-paraffinized by sequential immersions in xylene (bottles labeled #1, #2, and #3) for 10 min each, followed by graded ethanol washes (100%, 95%, and 70%) with 20 dips in each solution. Slides were rinsed in water before antigen retrieval using citrate buffer (cat# k035, Diagnostic Biosystems) in a sealed container for 30 min, followed by cooling for 30 min. Slides were then rinsed twice with PBS, and incubated for 15 min in Blocking Buffer to block endogenous peroxidase. After three washes with PBS, slides were incubated with primary antibody overnight at 4, and washed with PBS three times. Slides were then incubated with HRP conjugated secondary antibody for 1 h, followed by three PBS washes. Slides were air-dried and treated with 3,3’-diaminobenzidine (DAB) for 10 min, then rinsed with water and counterstained with hematoxylin for 10 min. Slides were sequentially immersed in water, 70%, 95%, and 100% ethanol for 10 min each, then mounted with Histomount or Permount for image analysis.

### Establishment of Huh7 VASN knockout cell lines

To establish a stable knockout of VASN in Huh7 and HepG2 cells, cells were first transfected with the Lenti-iCas9-Neo virus for 48 h. Cells were then passaged and selected using 4 μg/mL G418 (Cat#97064-358, VWR), until the control cell population reached 100% lethality. The selected viable cells were then transfected with VASN gRNA (forward sequence 5’-CACCGACGTGCCACCCGACACGGTG-3’, reverse sequence 5’-AAACCACCGTGTCGGGTGGCACGTC-3’) lentivirus to yield VASN knockout (KO) cells, or with control gRNA (a gift from Dr. Li Ma, MD Anderson Cancer Center) to yield the Control sRNA cells. Cells were then selected with puromycin (Abcam; 2 μg/ml for Huh7, 7 μg/ml for HepG2) until the control cell population exhibited complete lethality. The selected cells were treated with 4 μg/mL doxycycline (cat# AAJ6267106, ThermoFisher) for 72 h. The resulting VASN knockout cell lines were analyzed for knockout efficiency using Western blotting.

### Cell colony formation assay

Huh7 control gRNA or VASN knockout cells (1×10^3^), and HepG2 control gRNA or VASN knockout cells (2×10^3^) were seeded in 6-well plates and cultured for two weeks under standard conditions (37°C, 5% CO_2_, and 95% humidity), with a change of fresh, respective medium every 48 h. Cells were then fixed with methanol for 15 min at room temperature and stained with a 0.5% crystal violet solution for 10 min to visualize cellular morphology and adherence. Cells were then washed three times with distilled water to remove excess crystal violet, and the 6-well plates were air-dried before visualization and image acquisition using a high-resolution scanner (EPSON Perfection V500 PHOTO).

### *In vitro* wound healing assay

Cell migration was evaluated using a wound healing assay. Huh7 control gRNA or VASN knockout cells (3×10^5^) were seeded in a 12-well plate and allowed to form a confluent monolayer over a 24-h period. A wound was created by carefully scratching the bottom of the well using a sterile 200 μL pipette tip. Following wound creation, wells were washed with 500 μL of PBS to remove detached cells. Medium devoid of fetal bovine serum (FBS) but supplemented with 200 µg/mL bovine serum albumin (BSA) was introduced, and the cells were incubated for an additional 24 h. Subsequently, the plate was washed with 500 μL PBS and gently agitated for 30 seconds. Photos of regions surrounding the scratch were taken at the time of wound induction (0 h) and 48 h later, using a light microscope (Nikon).

### *In vivo* conditional knockout of VASN *via* doxycycline induction

Animal studies were carried out in compliance with federal and local institutional rules for the conduct of animal experiments. Two million Huh7 Control gRNA or Huh7 VASN KO cells were subcutaneously injected into the right shoulder of 6- to 10-week-old male or female NSG mice (n=10 to 20 in each group). On the day of cell inoculation, all mice were provided with drinking water containing 200 µg/mL of doxycycline, which continued until the experiment ended on day 25.

Tumor growth was monitored by measuring tumor dimensions using a Vernier caliper (Fisher scientific) twice a week, starting from the second week post-inoculation. From the third week onwards, the tumor size was measured every other day to ensure the tumor size did not exceed acceptable limits. Mice were sacrificed on day 25, and the harvested tumor volumes were calculated using the following equation:

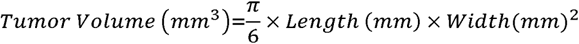

### RNA sequencing and data analysis

Total RNA was extracted from cell lines or tumor tissues using RNeasy Plus Mini Kit (cat# 74136, Qiagen). Subsequent mRNA library preparation and sequencing, and data analysis were done by Novogene using their standardized protocols. For non-directional libraries, the process included end repair, A-tailing, adapter ligation, size selection, amplification, and purification, while directional libraries involved additional steps such as USER enzyme digestion. The constructed libraries were checked for their quality with Qubit, quantified by real-time PCR, and analyzed for size distribution using the Bioanalyzer. These libraries were then pooled based on effective concentration and data amount before being sequenced on Illumina platforms.

The bioinformatics analysis pipeline comprised several stages, with robust methodologies that have been previously verified^27,28^. Initially, data quality control was performed to obtain clean reads by removing adapters, poly-N-containing reads, and low-quality reads. Subsequently, reads were mapped to the reference genome using Hisat2, and novel transcripts were predicted using StringTie. The quantification of gene expression levels was achieved with FeatureCounts, and differential expression analysis was conducted using DESeq2 or edgeR depending on the presence of biological replicates. GO and KEGG enrichment analyses were carried out, and Gene Set Enrichment Analysis (GSEA) was performed using predefined gene sets. SNP analysis was conducted with GATK, alternative splicing analysis with rMATS, and protein-protein interaction analysis based on the STRING database.

### Statistical analysis

Data were expressed as means ± standard error of the mean (SEM). Statistical comparisons between multiple groups were conducted using one-way analysis of variance (ANOVA) with GraphPad Prism software. Subsequent post-hoc analysis was performed using Tukey’s test to evaluate differences between individual groups. A p value of < 0.05 was considered to be statistically significant.

## Results

### NIC-PS exhibited similar potency and selectivity as NIC in HCC cell lines and primary human hepatocytes

To improve the poor water solubility and bioavailability of NIC, we developed a water improved prodrug, NIC-PS, synthesized, purified, and characterized by Acme Bioscience. To evaluate the impact of this structural modification on NIC’s anti-tumor activity, NIC, NIC-PS, and NEN (positive control) on the proliferation of HCC cell lines: Huh7, HepG2, and Hep3B. The three compounds are equipotent in HCC cell lines, with no significant different in IC_50_ values (Table S1), indicating that NIC-PS maintained the anti-tumor potency of NIC, and was equally potent as NEN. Additionally, NIC-PS was similarly non-cytotoxic to primary hepatocytes, suggesting its safety in normal liver cells.

### NIC pulled down vasorin from HCC cell lysates and bound to recombinant vasorin

To identify potential target proteins that might bind to NIC, we conducted a pull-down assay followed by mass spectrometry (Figure 1A). Coomassie Blue staining of SDS-PAGE gel (Fig. 1B) revealed potential protein bands of interest. Mass spectrometry identified these proteins, with vasorin (VASN) emerging as a top candidate with known biological functions. To validate this interaction, we performed a thermal shift assay using NEN (due to NIC’s limited solubility), confirming NEN’s binding to VASN with a dissociation constant (K_d_) of 6 µM (Fig. 1C).

**Figure 1.**
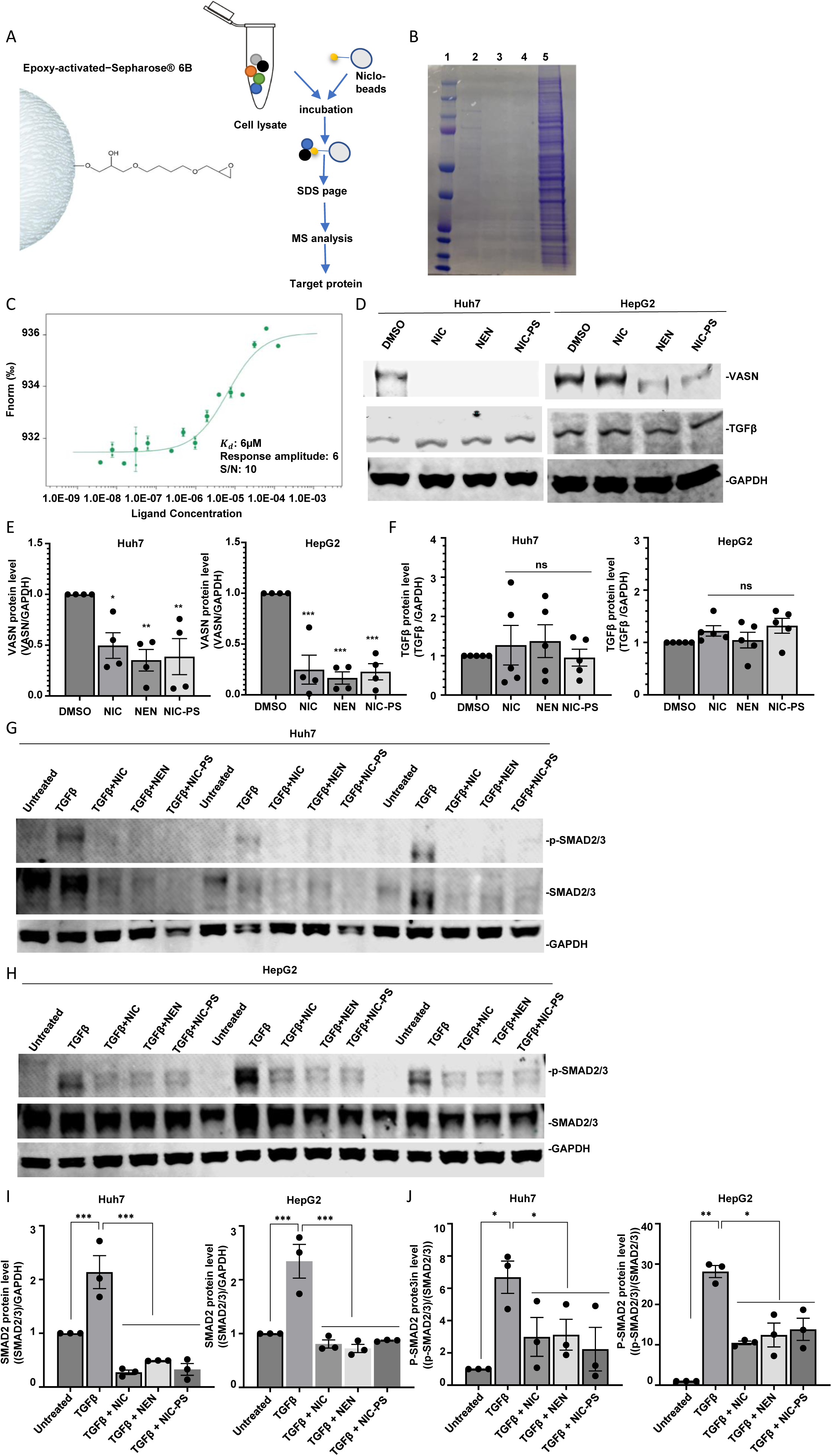
NIC bound to VASN and NIC-PS suppressed VASN and TGFβ signal pathway. **A)** Schematic of pull-down assay using NIC-resin beads to bind target proteins in Huh7 cell lysate. **B)** Coomassie blue staining of SDS-PAGE gel to analyze pulled down lysates. Lane 1: molecular weight marker, Lane 2: NIC-conjugated beads (incubated with 300 μg cell lysate), Lane 3: control beads (300 μg cell lysate), Lane 4: control beads (600 μg cell lysate), Lane 5: Huh7 cell lysate (300 μg). **C)** Thermal shift assay of NEN and VASN. **D)** VASN and TGFβ protein expression after treating Huh7 or HepG2 cells with 10x of the IC_50_ of NIC, NEN, or NIC-PS for 24 h. Quantified results for VASN and TGFβ are shown in **E and F)**, respectively. **G and H)** Duplicate experiments showing SMAD2/3 and p-SMAD2/3 protein levels after treating Huh7 **(G)** or HepG2 **(H)** cells with 10x of the IC_50_s of NIC, NEN, or NIC-PS for 24 h after 16 h fast without FBS. Quantified results for SMAD2/3 and p-SMAD2/3 are shown in **I and J**), respectively. Data are presented as mean ± SEM; significance levels are denoted as *, **, and *** for p-values < 0.05, 0.01, and 0.001, respectively, in comparison to the control group.

### NIC and NIC-PS suppressed VASN expression and TGFβ signaling pathway

To explore the effects of NIC and NIC-PS, HepG2 and Huh7 cells were treated, resulting in suppressed VASN protein levels (Fig. 1D & 1E), but not TGFβ protein levels (Fig. 1D and 1F). We also assessed the downstream of the TGFβ signaling pathway - total SMAD2/3 and their phosphorylation levels. NIC-PS reduced both total SMAD2/3 (Fig. 1G to 1I) and their phosphorylated forms (Fig. 1G, 1H and 1J). These findings suggest that NIC-PS effectively suppresses VASN and TGFβ signaling pathway. Although IL-33 has been reported to function within the TGFβ signaling loop^29^, we did not detect any changes (Fig. S1A-C).

### NIC-PS suppressed *in vivo* growth of orthotopic patient-derived xenograft of HCC

To assess whether NIC-PS retains the *in vivo* growth inhibitory effect, we first confirmed its bioavailability using a single dose pharmacokinetic study. Compared to NIC (40 mg/kg), NIC-PS (equivalent to 40 mg/kg of NIC based on molecular weight) showed an almost ten-fold increase in the area under the curve (AUC) of NIC levels in blood (Fig. 2A and Fig. S2A), demonstrating that NIC-PS is effectively cleaved to release NIC and enhances the NIC’s bioavailability. We hypothesized that NIC-PS may, therefore, increase tumor levels of NIC and its anti-tumor activity.

**Figure 2.**
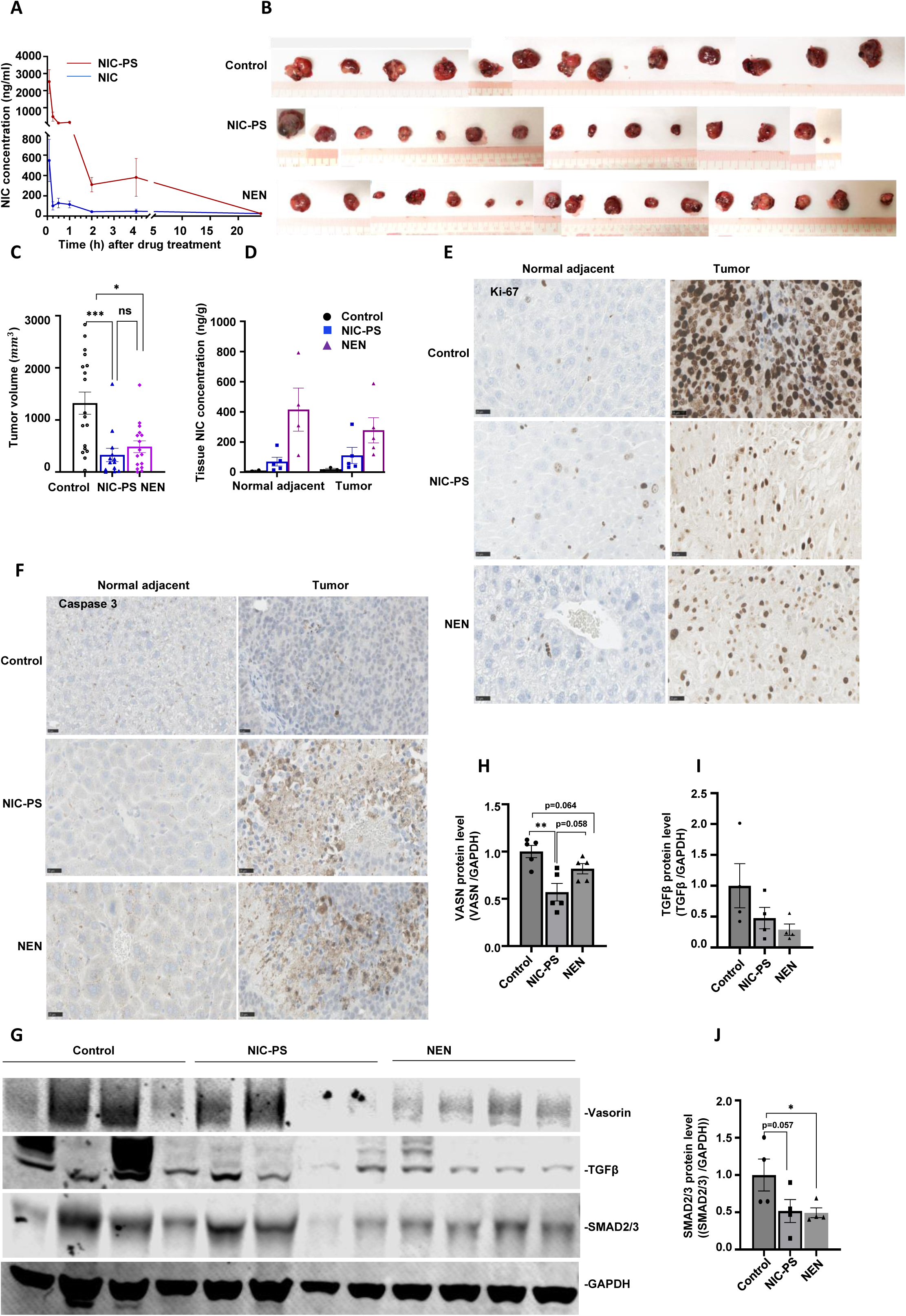
NIC-PS inhibited orthotopic HCC PDX growth. **A)** Blood NIC after a single oral dose of NIC (40 mg/kg) or NIC-PS (equivalent to 40 mg/kg of NIC) to non-tumor bearing mice. Plasma was collected from each group (n=3) at indicated time points and NIC levels were measured by LC-MS/MS. **B)** Harvested tumors and **C)** their tumor volumes from PDX-bearing mice after 4 weeks treatment with distilled water (untreated), NIC-PS or NEN for 4 weeks; (n=13 to 17). **D)** NIC levels in PDX tumor and normal adjacent liver (n=3 to 6). IHC staining of **(E)** Ki-67 and **(F)** Caspase-3 in PDX tumor and normal adjacent liver. G) VASN, TGFβ, SMAD2/3 protein levels in PDX tumor tissues after treatment with NIC-PS or NEN, and their quantification in **(H)**, **(I)**, and **(J)**, respectively. Data are presented as mean ± SEM; significance levels are denoted as *, **, and *** for p-values < 0.05, 0.01, and 0.001, respectively, compared to the untreated group.

In our *in vivo* efficacy study, we used NEN as a positive control, as previously reported ^5^. Based on molecular equivalents, the amount of NIC in the 100 mg/kg dose of NIC-PS was equivalent to ∼36% of NIC in 200 mg/kg dose of NEN; yet, both NIC-PS and NEN achieved significant PDX growth suppression, with tumor volume reductions of 75.13% for NIC-PS and 63.17% for NEN (Fig. 2B and 2C). Orthotopic PDX growth was monitored weekly by bioluminescence imaging (Fig. S2B). No detrimental effects were observed on body weight, and liver or kidney functions after the treatment period (Fig. S2C, Table S2). A second independent confirmed these findings, showing consistent tumor growth inhibition and similar safety outcomes on body weight and liver and kidney functions (Fig. S2D, 2E, and S2F, Table S3).

We measured NIC levels in harvested PDX tissues and adjacent normal livers after 4-week treatment. In the NEN-treated group, NIC levels in the normal adjacent livers were ∼1.3 times higher than in the PDX tumors (Fig. 2D). However, in the NIC-PS-treated group, NIC levels in the PDX tumors were ∼1.5 times higher than in the normal adjacent livers (Fig 2D), suggesting enhanced accumulation of NIC in the target tumor. Additionally, NIC levels in adjacent liver were ∼4 times higher with NEN than with NIC-PS. Both the preferential accumulation of NIC in the PDX, and the lower levels of NIC in the normal liver suggest that the NIC-PS is less toxic to the normal liver, than NEN. Even though PDX levels of NIC (when was NIC-PS administered) was lower than when NEN was given (consistent with the lower NIC molecular equivalent), NIC-PS achieved comparable tumor inhibition at a lower dose. These observations highlight the superior performance and safety profile of NIC-PS. Indeed, neither NEN nor NIC-PS affected the mice body weight (Fig. S2C and S2F) or kidney and liver functions (Tables S2 and S3) as mentioned above. Consistent with previous report on NIC^20,30^, NIC-PS decreased blood sugar levels (from 246.9 ± 49.7 mg/dL on Day 0 to 149.0 ± 28.7 mg/dL on Day 28).

At the biochemical level, both NIC-PS and NEN inhibited Ki-67 expression (Fig. 2E), with NIC-PS exhibiting greater inhibition, suggesting a stronger anti-proliferative effect. Both compounds similarly elevated caspase-3 levels (Fig. 2F). Consistent with *in vitro* data, NIC-PS and NEN treatment in PDX-bearing mice decreased VASN, TGFβ, and SMAD2/3 protein levels in the tumors, NIC-PS suppressed the VASN protein level by 58.4%, TGFβ by 52.4%, and SMAD2/3 by 48.3%, compared to the untreated group (Fig. 2G-J). Similar to *in vitro* data, no significant drug-induced changes in IL33 protein levels in the harvested tumors (Fig. S3A and S3B) or in mouse plasma (Fig. S3C), nor in plasma TGFβ (Fig. S3D). However, VASN plasma levels increased (Fig. S3E).

### Suppression of VASN inhibited HCC cell proliferation and migration

Based on our findings that NIC binds to and suppresses VASN, which may be the novel target of NIC, we explored the functions of VASN in HCC cells. We first investigated VASN expression in the immortalized non-tumor human liver cell line (THLE2) and a panel of HCC cell lines, and observed negligible VASN level in the THLE2 cells, but distinct VASN expression in all HCC cells lines, with the highest levels in Huh7 cells (Fig. 3A and B). We then suppressed VASN expression in Huh7 and HepG2 cells, achieving a 51-98% knockdown efficiency (Fig. 3C and 3D). VASN knockout in both Huh7 and HepG2 significantly reduced colony formation and cell migration (Fig. 3E and F).

**Figure 3.**
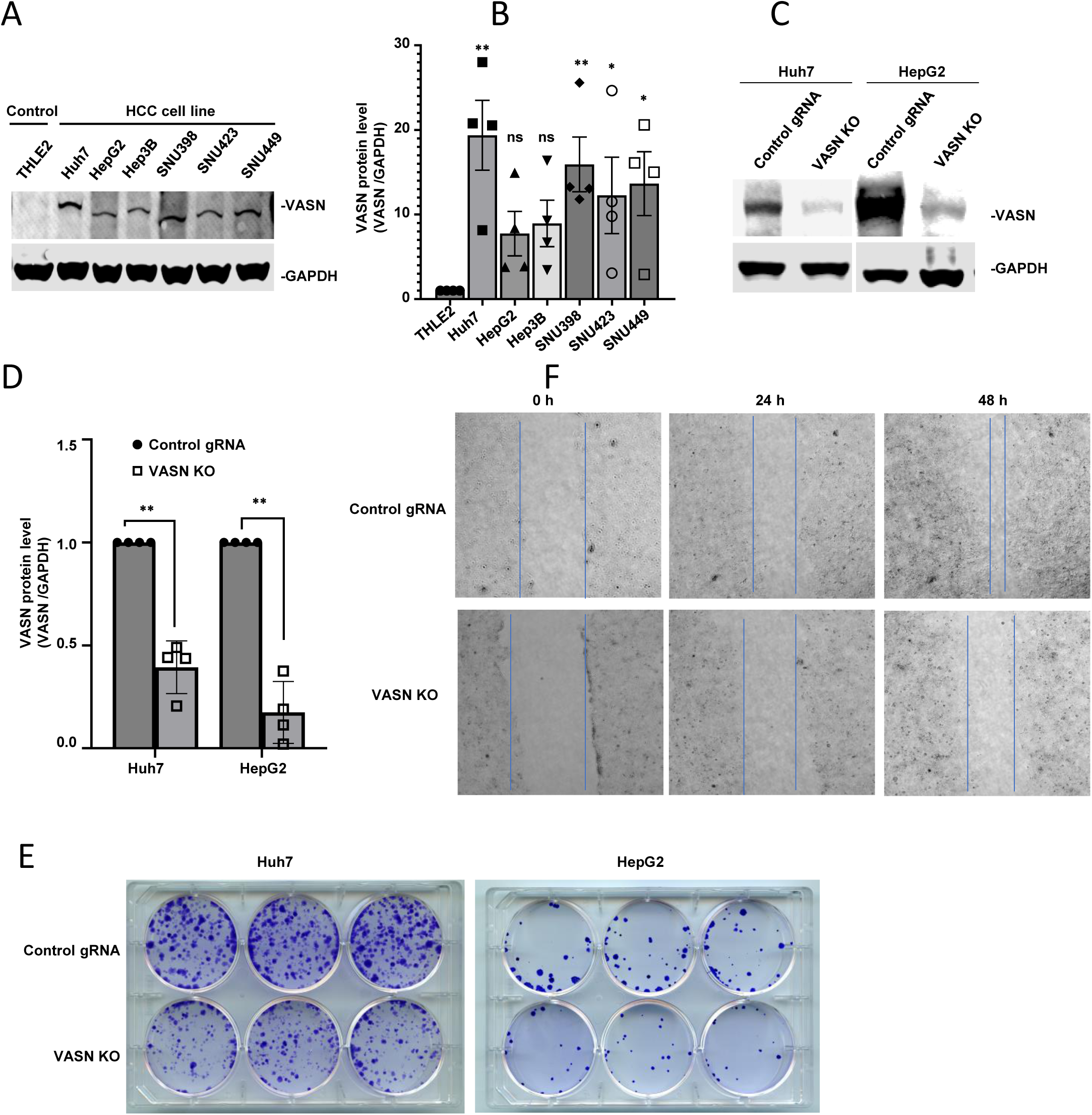
VASN knockout in HCC cells inhibited colony formation and cell migration. **A)** VASN protein level in immortalized non-tumor liver cells (THLE2) and a panel of HCC cell lines, and their **B)** quantified results. **C)** Stable VASN knockout in Huh7 and HepG2 cells, and quantified in **D)**. **E)** colony formation and **F)** cell migration abilities of Huh7 cells. Data are presented as mean ± SEM; significance levels are denoted as * and **, for p-values < 0.05 and 0.01, respectively, in comparison to THLE2 cells.

Collectively, these findings support the role of VASN in HCC cell proliferation and migration.

### VASN knockout suppressed *in vivo* growth of subcutaneous Huh7 xenografts

To extend our observations, we suppressed VASN through a doxycycline-inducible system in Huh7 cells (Huh7-VASN KO), which were then subcutaneously injected into NSG mice. Doxycycline was administered in the drinking water for three weeks to induce VASN suppression, leading to a significant reduction in xenograft growth compared to control Huh7 cells (Huh7 Control gRNA), achieving approximately 50% tumor volume suppression (Fig. 4A-C). VASN protein was nearly absent in the harvested Huh7-VASN KO xenografts (Fig. 4D), confirming an 87.5% knockdown efficiency (Figure 4E). Additionally, Ki-67 levels decreased, while Caspase-3 levels increased in the Huh7-VASN KO xenografts compared to the control (Fig. 4F and G). Consistent with *in vitro* data, there are no significant changes in TGFβ (Fig. 4H and I) or IL33 (Fig. S4A and S4B) protein levels in the xenografts.

**Figure 4.**
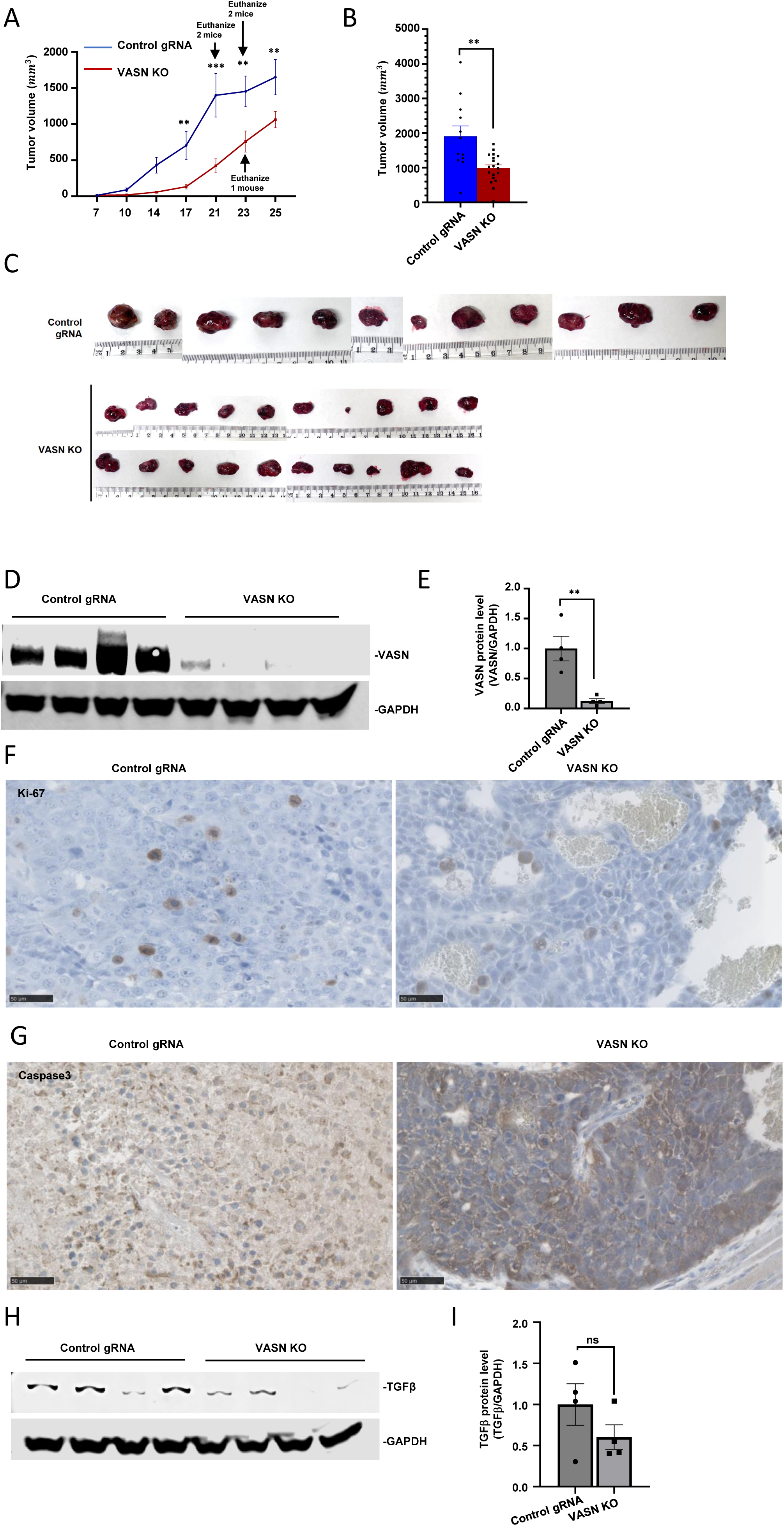
VASN knockout inhibited Huh7 xenograft growth. **A)** Tumor growth curves in Huh7 control gRNA and VASN KO cells over 25 days, with VASN knockout induced by 200 μg/mL doxycycline drinking water. **B)** Tumor volume calculations and **C)** images of harvested tumors (n=12 to 20 per group). **D)** VASN knockout efficiency in tumors and **E)** quantification. **F)** Ki-67 and **G)** Caspase-3 staining in harvested xenografts. **H)** TGFβ protein levels in harvested xenografts, and their I) quantified results. Data are presented as mean ± SEM; significance levels are denoted as ** and *** for p-values < 0.01 and 0.001, respectively, in comparison to the control.

### NIC-PS treatment or VASN knockout increased susceptibility of HCC cells to anti-PD-L1 and sorafenib

To evaluate NIC-PS as a complementary therapy for HCC, we assessed its potential to enhance the effects of sorafenib and anti-PD-L1 immunotherapy. NIC, NEN, and NIC-PS all reduced PD-L1 mRNA and protein levels in Huh7 and HepG2 cells (Fig. 5A-C).

**Figure 5.**
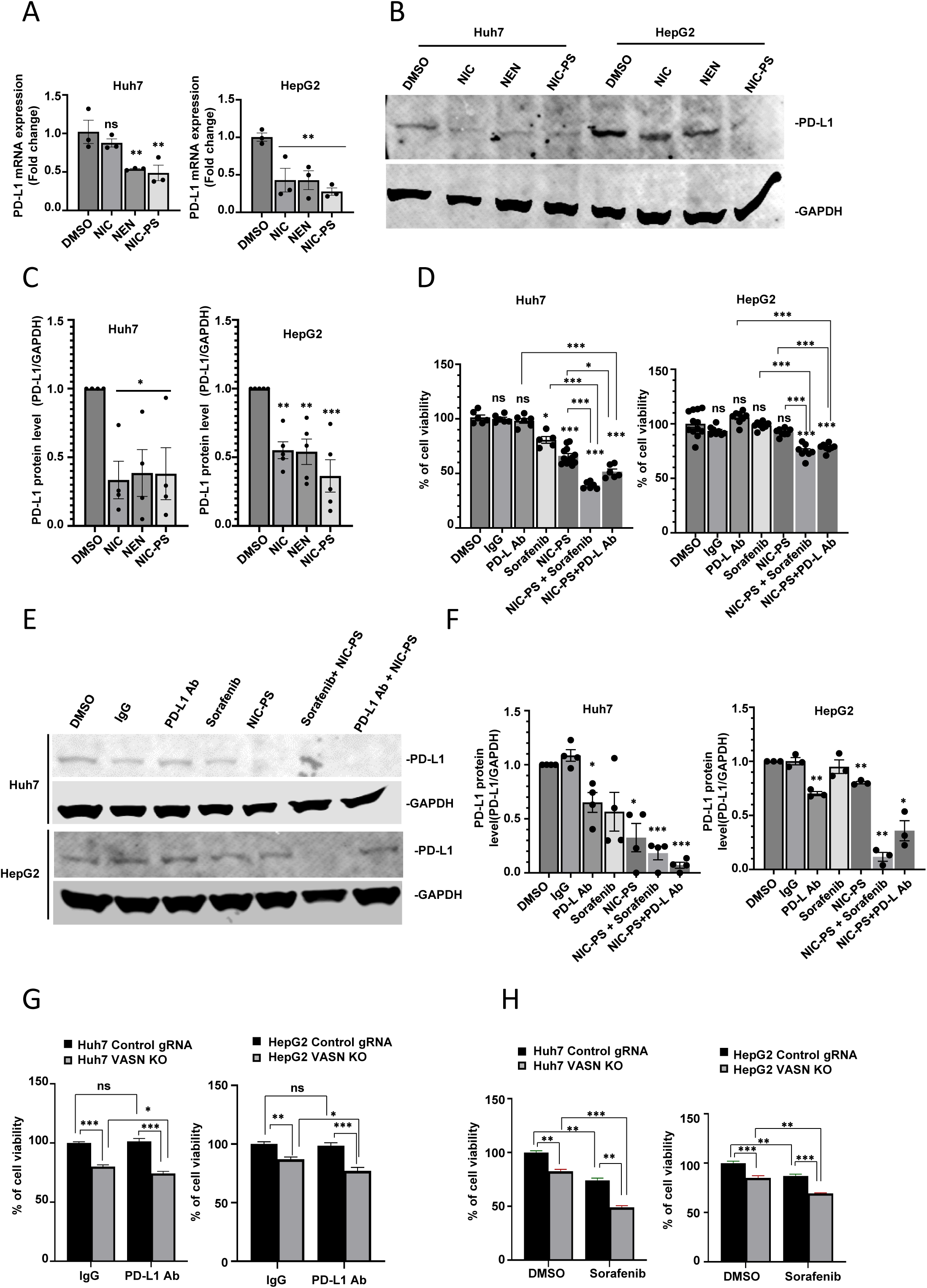
NIC-PS and VASN knockout decreased PD-L1 and enhanced HCC susceptibility to sorafenib and anti-PD-L1 treatment. **A)** PD-L1 mRNA expression in Huh7 and HepG2 cells after 6 hours of treatment with 10x IC50 of NIC, NEN, or NIC-PS. **B)** PD-L1 protein levels in Huh7 and HepG2 cells after 72 hours of treatment with IC50 of NIC, NEN, or NIC-PS. **C)** Quantification of PD-L1 protein levels. **D)** Enhanced sensitivity of Huh7 and HepG2 cells to anti-PD-L1 (10 μg/mL) or sorafenib (5 μM) when co-treated with NIC-PS (1.36 μM in Huh7 or 2.2 μM in HepG2) for 72 hours. **E** and **F)** Co-treatment of NIC-PS with anti-PD-L1 or sorafenib further reduced PD-L1 protein levels. **G** and **H)** VASN KO cells in Huh7 and HepG2 showed increased sensitivity to co-treatment with anti-PD-L1 (10 μg/mL) or sorafenib (5 μM). Data are presented as mean ± SEM, significance levels are denoted as *, **, and *** for p-values < 0.05, 0.01, and 0.001, respectively, in comparison to the control.

Next, Huh7 and HepG2 cells were treated with NIC-PS alone, in combination with sorafenib, or with an anti-PD-L1 antibody. Co-treatments with NIC-PS and either sorafenib or anti-PD-L1 significantly enhanced the sensitivity of these cells compared to treatment with sorafenib or anti-PD-L1 alone (Fig. 5D), accompanied by further reductions in PD-L1 protein levels (Fig. 5E-F). VASN KO cells showed increased sensitivity to both anti-PD-L1 (Fig. 5G) and sorafenib (Fig. 5H), supporting that NIC-PS enhances susceptibility through VASN inhibition. These results suggest NIC-PS could be clinically useful in combination therapy with sorafenib or anti-PD-L1, likely by inhibiting VASN and TGFβ activity.

### NIC-PS regulated HCC growth *via* multiple mechanisms

To further investigate the mechanisms underlying NIC-PS-mediated suppression of HCC growth, we analyzed differentially expressed (DE) genes in untreated PDX versus NIC-PS-treated PDX. We then compared these with DE genes in Huh7-VASN KO xenografts versus Huh7 Control gRNA xenografts and in untreated Huh7 cells versus NIC-treated cells. Comparative analysis identified several signaling pathways affected by NIC and VASN KO (Fig. 6A and B). KEGG and GO analyses revealed significant pathways in the NIC-PS-treated versus untreated groups, including oxidative phosphorylation, ribonucleoside monophosphate metabolism, purine ribonucleotide metabolism, and mitochondrial inner membrane protein complexes (Fig. 6C-E). Specifically, Huh7 VASN KO was associated with the complement and coagulation cascades, staphylococcus aureus infection, TGFβ signaling, and PI3K-Akt signaling (Fig. 6F-H). These findings align with known NIC mechanisms^7,31^ and our observations on its impact on the TGFβ pathway.

**Figure 6.**
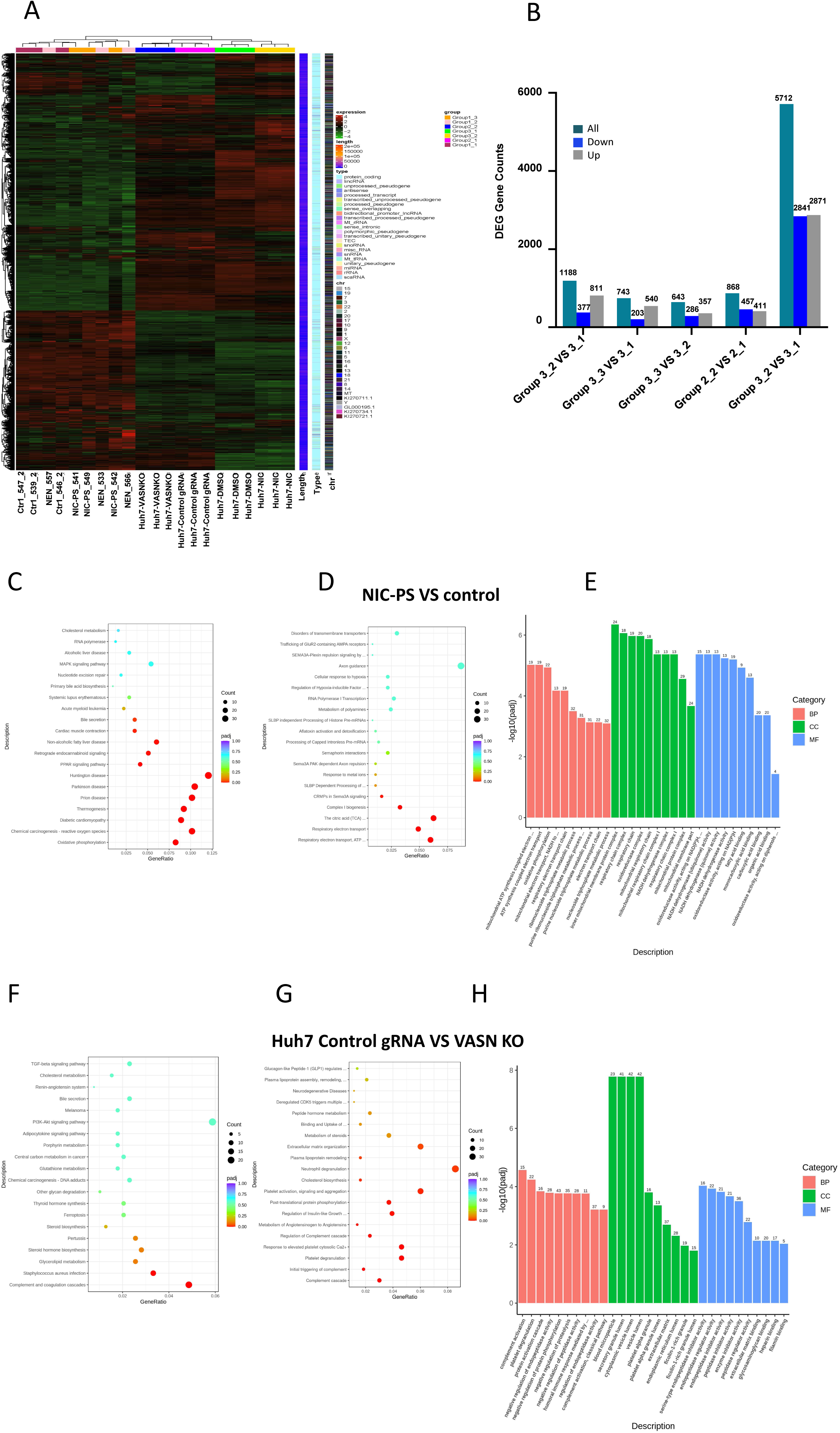
NIC-PS and VASN knockout affect HCC growth via multiple signaling pathways. **A)** Heatmap showing the differentially expressed genes based on RNA sequencing of PDX-control (Group 1-1), PDX-NEN (Group1-2), PDX-NIC-PS (Group 1-3), Huh7 control gRNA xenograft (Group 2-1), Huh7 VASN KO xenograft (Group2-2), Huh7 parental cell xenograft (Group 3-1), Huh7 xenograft treated with NIC (Group3-2). **B)** The number of differentially expressed gene counts in each comparison group (shown on x-axis). **C** to **E)** GO and KEGG analysis data showing major signaling pathways affected after treating Huh7 cells with NIC-PS. **F** to **H)** The major signaling pathways affected after VASN knockout in Huh7 cells. BP – biological process, CC – cellular component, MF – molecular function.

## Discussion

The success rate of HCC treatment remains low despite advances in molecular understanding and clinical trials with newer receptor tyrosine kinase inhibitors and immunotherapeutic modalities^4^. Our study introduces a novel approach to repurpose NIC for HCC therapy by addressing its poor water solubility and bioavailability.

Prior studies, including our own, highlighted NIC’s poor solubility and bioavailability as the key limitations impeding its effectiveness against solid tumors in clinical application^32^. To overcome these challenges, we developed NIC-PS, and evaluated its bioavailability, *in vivo* anti-tumor efficacy in two orthotopic HCC-PDX mouse models, and mechanisms of action.

Pharmacokinetic studies confirmed that NIC-PS converts effectively to NIC, resulting in higher and prolonged plasma levels compared to oral NIC. This higher bioavailability, evidenced by increased NIC levels in PDX tissues and preferential accumulation in tumors, correlated with improved anti-tumor efficacy.

Mechanistically, we identified NIC’s binding to VASN in Huh7 cells, this is the novel discovery which has not been reported yet. VASN is known to stimulate angiogenesis and tumor progression ^33^ and binds TGFβ to block its activity ^34^. Our data show that NIC, NEN, and NIC-PS reduce VASN levels and TGFβ activity, as indicated by decreased SMAD2/3 levels and phosphorylation. SMAD2/3 are key effectors in the TGFβ signaling pathway, and their phosphorylation is crucial for TGFβ activation. Notably, SMAD2/3 was also reported to be involved in the function of NIC in alleviating pulmonary fibrosis and inhibiting epithelial-mesenchymal transition in BRAF/ NRAS mutated metastatic melanoma^35,36^. Increases in soluble VASN in the plasma of mice treated with NIC-PS or NEN suggest a mechanism involving suppression of TGFβ signaling; since soluble VASN was reported to inhibit TGFβ signaling^34^.

We further demonstrated VASN’s role in HCC by knocking out VASN in Huh7 and HepG2 cells. Both NIC and NIC-PS treatments reduced VASN levels and associated TGFβ activity, aligning with the effects seen in VASN knockout cells. This suggests that HCC growth suppression by NIC and NIC-PS may partially involve the VASN-TGFβ axis, warranting further investigation.

We observed elevated VASN protein expression in HCC cells while low VASN expression in normal liver cells, suggesting that VASN may be a safe therapeutic target with minimal toxicity to major organs. Transcriptomic analysis of Huh7 VASN knockout cells revealed suppressed TGFβ signaling, supporting the therapeutic potential of VASN inhibition. VASN’s involvement in HCC is further supported by its reported roles in enhancing Notch signaling under hypoxic conditions ^37^ and promoting HCC growth *via* STAT3 signaling ^38^. Our data, along with established functions of VASN in HCC, underscore its potential as a therapeutic target, with transcriptomic data revealing its multifaceted involvement in HCC cells.

We previously reported that NIC and sorafenib co-treatment caused a greater inhibitory effect on HCC cell growth *in vitro* and *in vivo*, than either agent given alone^5^. Here, NIC-PS co-treatment with sorafenib achieved similar outcomes in vitro, as did sorafenib treatment of VASN knockout cell lines. Similarly, NIC-PS and VASN knockout enhanced sensitivity to anti-PD-L1 antibody treatment. These findings support NIC-PS as a potentially valuable therapeutic agent, either alone or in combination with other established therapies. Additionally, NIC-PS decreased PD-L1 protein levels, suggesting it may act via PD-L1 inhibition. PD-L1 is a major checkpoint regulator, when inhibited, can prevent tumor cells from evading T cell killing, which is crucial for overcoming resistance to immunotherapy ^39^. Thus, targeting PD-L1 is a promising cancer treatment strategy ^40^. Our study revealed a novel mechanism of NIC, suggesting that it effectively inhibits the TGFβ signaling through VASN suppression and enhances HCC cell susceptibility to PD-L1 blockade.

TGFβ, elevated in various tumors, including colorectal, breast, and lung cancers, promotes tumor growth and invasiveness while also regulating immune responses. It aids tumor cells in evading immune surveillance, leading to resistance against cancer immunotherapy^41,42^. Indeed, accumulating evidence suggest that TGFβ can be classified as the checkpoint’s inhibitor^41,42^. In HCC, TGFβ plays a dual role in tumor biology, suppressing T cell growth and mediating immune escape ^43^. Perez *et al* demonstrated that the synergistic effects of blocking TGFβ with anti-CTLA4 and anti-PD1 therapies effectively suppressed bone metastasis in murine prostate cancer models^44^. Dodagatta-Marri *et al^45^* and Laine *et al^46^* reported that TGFβ activation through integrin αVβ8 leads to immune escape. These studies underscore the importance of inhibiting TGFβ for effective cancer therapy, particularly in HCC where it impacts inflammation, fibrogenesis, and microenvironment modulation^47^. Since TGFβ has been shown to reduce the efficacy of PD-L1 blockade ^48^, the suppression of TGFβ by NIC-PS and VASN KO may explain the increased susceptibility of HCC cells to PD-L1 blockade and sorafenib. Our results align with earlier reports that NIC ^49^ and other anthelmintics^50^ enhance anti-tumor responses to PD-1/PD-L1 blockade, and offer novel insights into the mechanisms.

Our study did not include **1)** biochemical experimentation on NIC’s binding to VASN or its effects on VASN expression and downstream activities due to the lack of a validated VASN crystal structure; **2)** Molecular docking studies may provide valuable NIC’s binding sites and interaction mechanisms with VASN. Future work will focus on these studies, refining dosages and regimens to optimize therapeutic outcomes and safety. Additionally, further mechanistic studies on NIC-PS’s impact on the tumor microenvironment and host immune system are needed to support its inclusion in HCC treatment.

In conclusion, we have developed NIC-PS, a water-soluble, orally bioavailable NIC prodrug that overcomes NIC’s clinical limitations and enhances anti-tumor efficacy. Our study is the first to identify VASN as a key NIC binding target in HCC cells and demonstrated that NIC-PS effectively inhibits the TGFβ pathway. Additionally, the enhanced efficacy of NIC-PS in combination with sorafenib and anti-PD-L1 therapies highlights its potential as a promising new therapeutic option for HCC.

## Acknowledgements

The authors would like to acknowledge the Stanford University Cell Sciences Imaging Core Facility (RRID:SCR_017787) for providing training and equipment access. We also thank Dr. Faiz Ahamad at Stanford University for assistance in animal studies, and Dr. Hongqi Teng at MD Anderson Cancer Center for technical suggestions. Dr. Danica Chen at University of California, Berkeley for providing Lenti-9Cas9-Neo plasmid.

## Authors contributions

Designed study: M.T., X.C., M.C. Participated in/executed trial/study: M.T., W.Y., Y.L., L.H., and X.C. Data collection/acquisition: M.T., W.Y., Y.L., and L.H. Data analysis: M.T., W.Y., and Y.L. Data interpretation: M.T., and M.C. Manuscript writing: M.T., W.Y., Y.L., and M.C. Supervision of study: M.C., and S.S. Approval of final submitted version: M.T., W.Y., Y.L., L.H., X.C., M.C., and S.S.

## Ethics approval and consent to participate

All animal studies were performed strictly according to the guidelines and rules concerning laboratory animal care, and were approved by the Institutional Animal Care and Use Committee (IACUC) at Stanford University (Protocol number: APLAC-20167). HCC tissues were collected from HCC patients who had undergone liver resection as part of their treatment. The protocol was approved by the Institutional Review Board at Stanford University, and informed consent was obtained from patients prior to the procedure.

## Consent for publication

All authors are consent for publication.

## Data availability

All data are available in the main text or the supplementary materials. NCBI RNA-sequencing raw data BioSample accession number: SAMN41537818, SAMN41537819, SAMN41537838; SAMN41537841, AAMN41537842; SAMN41537843, SAMN41537844.Materials and reagents utilized in this research can be obtained through the corresponding author, and specific conditions for access or collaboration can be discussed.

---

**Table S1.** IC_50_ values of NIC, NEN, and NIC-PS in HCC cell lines and in human primary hepatocytes.

|  | IC<sub>50</sub> (μM) (Mean ± SEM) |  |  |  |
| --- | --- | --- | --- | --- |
|  | Huh7 | HepG2 | Hep3B | Primary Human Hepatocytes |
| **NIC** | **0.79 ± 0.079** | **1.19 ± 0.066** | **0.49 ± 0.033** | **4.15 ~ 4.70** |
| **NEN** | **0.79 ± 0.088** | **1.06 ± 0.045** | **0.45 ± 0.002** | **3.30 ~ 7.76** |
| **NIC-PS** | **1.36 ± 0.15** | **2.24 ± 0.12** | **0.71 ± 0.081** | **5.41 ~ 6.74** |

**Table S2.** Biochemical analysis of blood at NIC-PS treatment endpoint in orthotopic HCC PDX mice.

|  | Mean ± SEM |  |  |
| --- | --- | --- | --- |
|  | Control | NIC-PS (100 mg/Kg) | NEN (200 mg/Kg) |
| WBC | 1.44 ± 0.24 | 2.2 ± 0.26 | 2.21 ± 0.16 |
| RBC | 7.3 ± 0.32 | 8.59 ± 0.05 | 7.97 ± 0.36 |
| HGB | 10.78 ± 0.49 | 12.78 ± 0.17 | 11.88 ± 0.63 |
| HCT | 35.66 ± 1.26 | 40.34 ± 0.58 | 37.44 ± 1.89 |
| MCV | 48.92 ± 0.57 | 46.98 ± 0.44 | 46.92 ± 0.58 |
| MCH | 14.78 ± 0.21 | 14.88 ± 0.17 | 14.86 ± 0.2 |
| MCHC | 30.22 ± 0.6 | 31.68 ± 0.26 | 31.7 ± 0.16 |
| Platelet Count | 2073 ± 251.44 | 1874 ± 115.11 | 1856 ± 146.91 |
| RDW | 16.44 ± 0.47 | 16.96 ± 0.13 | 16.6 ± 0.18 |
| PDW | 6.54 ± 0.08 | 6.46 ± 0.05 | 6.52 ± 0.25 |
| MPV | 6.22 ± 0.09 | 6.38 ± 0.08 | 6.44 ± 0.22 |
| P-lcr | 4.64 ± 0.31 | 4.28 ± 0.44 | 4.04 ± 1.36 |
| PCT | 1.08 ± 0.17 | 0.96 ± 0.06 | 0.99 ± 0.04 |
| Reticulocyte count | 4.84 ± 1.06 | 3.7 ± 0.44 | 4.48 ± 0.88 |
| IRF | 52.82 ± 6.17 | 54.98 ± 2.46 | 57.84 ± 2.42 |
| IFR | 47.18 ± 6.17 | 45.02 ± 2.46 | 42.16 ± 2.42 |
| MFR | 13.72 ± 1.03 | 13.04 ± 0.46 | 12.24 ± 0.66 |
| Reticulocyte Abs | 343610.4 ± 64028.09 | 317928.2 ± 38487.32 | 345706.2 ± 48280.68 |
| Platelet Estimate | Adequate | Adequate | Adequate |
| Neutrophils % | 88.4 ± 2.84 | 81.6 ± 2.46 | 89.6 ± 2.16 |
| Lymphocytes % | 6.8 ± 2.4 | 13.6 ± 1.94 | 7.6 ± 1.72 |
| Monocytes % | 4.8 ± 0.58 | 4.4 ± 0.93 | 2.4 ± 0.75 |
| Eosinophils % | 0 ± 0 | 0.4 ± 0.24 | 0.4 ± 0.24 |
| Basophils % | 0 ± 0 | 0 ± 0 | 0 ± 0 |
| RBC Morphology | Normal | Normal | Normal |
| Neutrophil Absolute | 1282 ± 215.37 | 1783.6 ± 197.33 | 1987.8 ± 193.42 |
| Lymphocytes Absolute | 93.6 ± 36.48 | 309.2 ± 73.38* | 159.6 ± 32.46 |
| Monocytes Absolute | 68.6 ± 15.35 | 97 ± 21.46 | 50.4 ± 16.6 |
| Eosinophils Absolute | 0 ± 0 | 6.6 ± 4.07 | 8.2 ± 5.04 |
| Basophil Absolute | 0 ± 0 | 0 ± 0 | 0 ± 0 |
| AST U/L | **615.4 ± 133.87** | **243.6 ± 82.07\*** | **313.4 ± 193.7** |
| ALT U/L | **112.2 ± 24.9** | **58.4 ± 9.52\*** | **60.8 ± 26.61** |
| GGT U/L | **37 ± 5.77** | **15.2 ± 6.46** | **14.2 ± 6.42\*** |
| Total Bilirubin mg/dL | 0.16 ± 0.04 | 0.18 ± 0.02 | 0.2 ± 0.05 |
| Direct Bilirubin mg/dL | — | 0.03 ± 0.03 | 0.08 ± 0.03 |
| Indirect Bilirubin mg/dL | — | 0.13 ± 0.03 | 0.15 ± 0.05 |
| BUN mg/dL | 31.4 ± 1.08 | 28 ± 1.05 | 46.4 ± 17.03 |
| Creatinine mg/dL | 0.43 ± 0.04 | 0.4 ± 0.03 | 0.56 ± 0.16 |
| Phosphorus mg/dL | 13.26 ± 1.87 | 9.4 ± 0.73 | 12.44 ± 2.54 |
| T. protein g/dL | 7.3 ± 0.58 | 8.2 ± 0.26 | 8.08 ± 0.7 |
| Albumin g/dL | 4.48 ± 0.31 | 4.92 ± 0.14 | 4.72 ± 0.26 |
| Globulin | 2.82 ± 0.28 | 3.28 ± 0.14 | 3.36 ± 0.45 |
| Cholesterol mg/dL | 211.6 ± 28.22 | 255.8 ± 16.96 | 247.2 ± 37.27 |
| Triglycerides mg/dL | 153.2 ± 43.11 | 256.8 ± 90.8 | 206.6 ± 56.28 |
| HDL mg/dL | 193.4 ± 27.17 | 240.6 ± 12.21 | 235 ± 33.19 |
| LDL mg/dL | 24.2 ± 3.31 | 15.6 ± 2.06 | **13.2 ± 3.48\*** |
| Uric Acid mg/dL | 0.92 ± 0.07 | 0.96 ± 0.2 | 1.04 ± 0.14 |

WBC: white blood cell, RBC: red blood cell, HGB: hemoglobin, HCT: Hematocrit, MCV: mean corpuscular volume, MCH: mean corpuscular hemoglobin , MCHC: mean corpuscular hemoglobin concentration, RDW: red cell distribution width, PDW: platelet distribution width, MPV: mean platelet volume, p-lcr: platelet-large cell ratio , PCT: procalcitonin, IRF: immature reticulocyte fraction, IFR: immature reticulocyte fraction, MFR: medium fluorescence reticulocytes, AST: Aspartate Transferase, ALT: Alanine Aminotransferase , GGT: Gamma-glutamyl Transferase, BUN: Blood urea nitrogen. Data are presented as mean ± SEM; \* denotes significance at p-values < 0.05, in comparison to the control group.

**Table S3.**
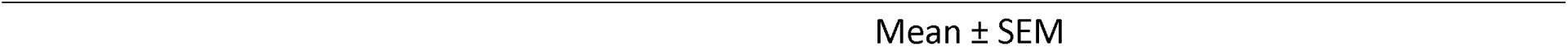
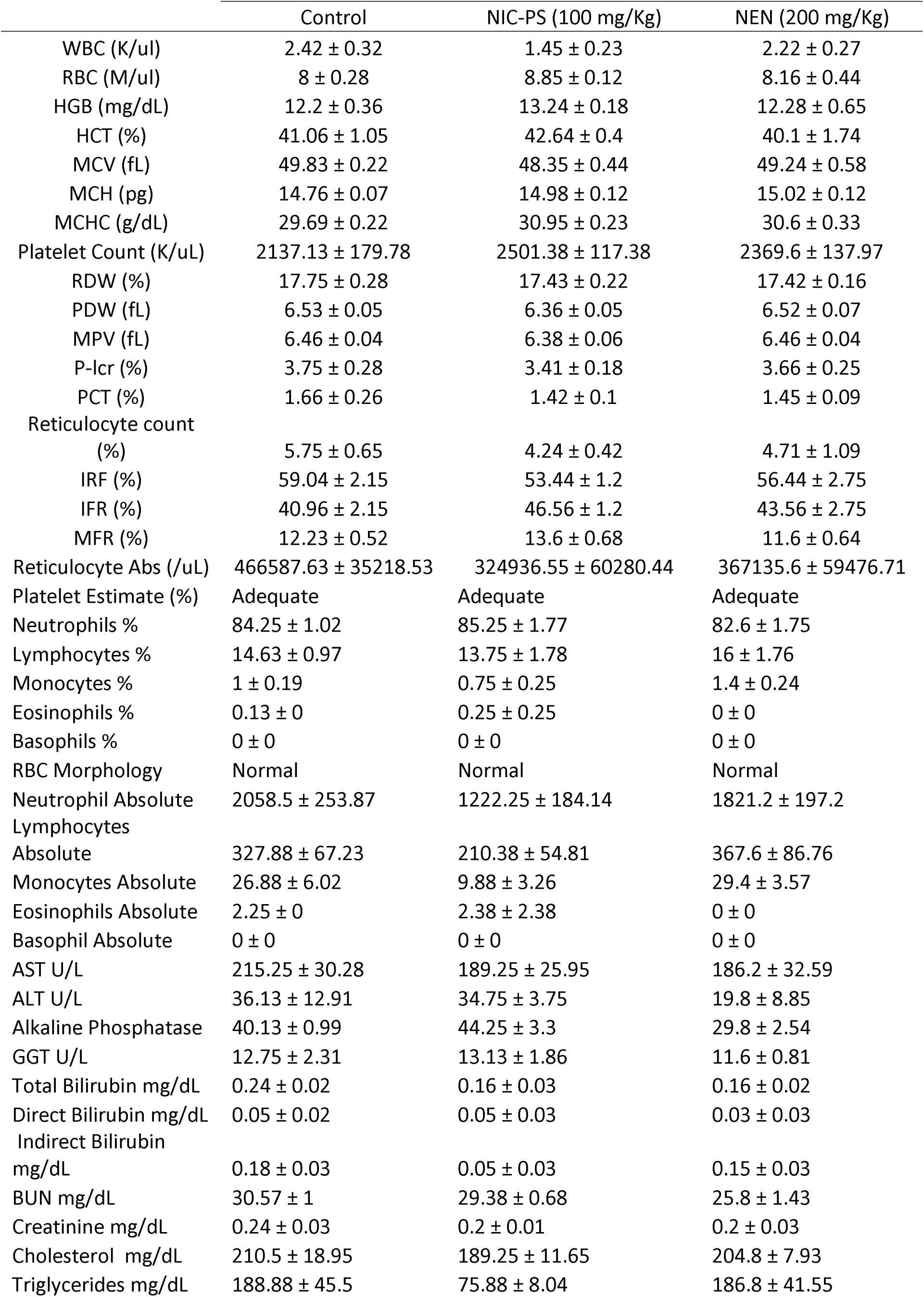
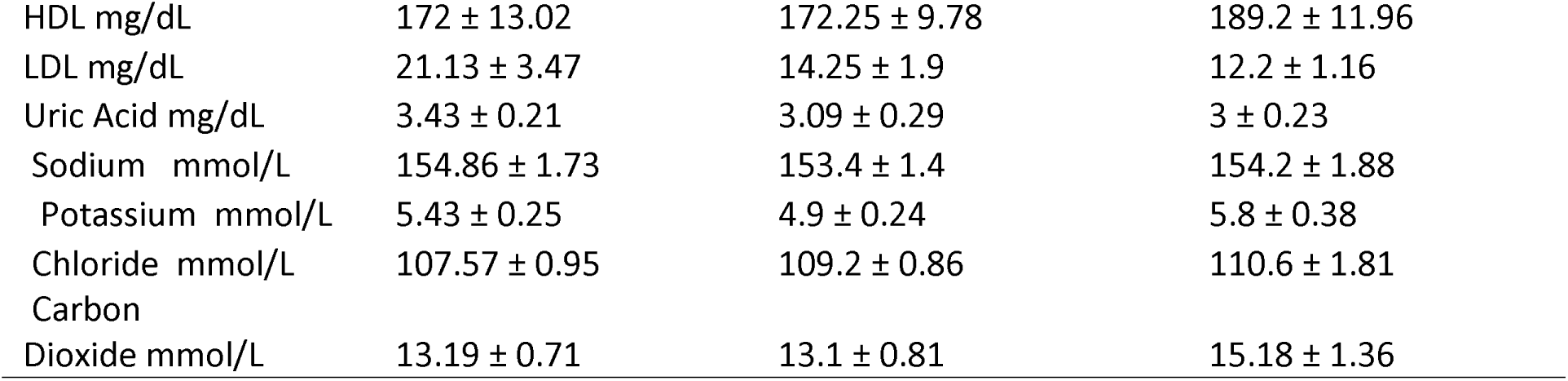
Biochemical analysis of blood at NIC-PS treatment endpoint in orthotopic HCC PDX mice in a second independent experiment.

|  | Control | NIC-PS (100 mg/Kg) | NEN (200 mg/Kg) |
| --- | --- | --- | --- |
| WBC (K/uL) | 2.42 ± 0.32 | 1.45 ± 0.23 | 2.22 ± 0.27 |
| RBC (M/uL) | 8 ± 0.28 | 8.85 ± 0.12 | 8.16 ± 0.44 |
| HGB (mg/dL) | 12.2 ± 0.36 | 13.24 ± 0.18 | 12.28 ± 0.65 |
| HCT (%) | 41.06 ± 1.05 | 42.64 ± 0.4 | 40.1 ± 1.74 |
| MCV (fL) | 49.83 ± 0.22 | 48.35 ± 0.44 | 49.24 ± 0.58 |
| MCH (pg) | 14.76 ± 0.07 | 14.98 ± 0.12 | 15.02 ± 0.12 |
| MCHC (g/dL) | 29.69 ± 0.22 | 30.95 ± 0.23 | 30.6 ± 0.33 |
| Platelet Count (K/uL) | 2137.13 ± 179.78 | 2501.38 ± 117.38 | 2369.6 ± 137.97 |
| RDW (%) | 17.75 ± 0.28 | 17.43 ± 0.22 | 17.42 ± 0.16 |
| PDW (fL) | 6.53 ± 0.05 | 6.36 ± 0.05 | 6.52 ± 0.07 |
| MPV (fL) | 6.46 ± 0.04 | 6.38 ± 0.06 | 6.46 ± 0.04 |
| P-lcr (%) | 3.75 ± 0.28 | 3.41 ± 0.18 | 3.66 ± 0.25 |
| PCT (%) | 1.66 ± 0.26 | 1.42 ± 0.1 | 1.45 ± 0.09 |
| Reticulocyte count (%) | 5.75 ± 0.65 | 4.24 ± 0.42 | 4.71 ± 1.09 |
| IRF (%) | 59.04 ± 2.15 | 53.44 ± 1.2 | 56.44 ± 2.75 |
| IFR (%) | 40.96 ± 2.15 | 46.56 ± 1.2 | 43.56 ± 2.75 |
| MFR (%) | 12.23 ± 0.52 | 13.6 ± 0.68 | 11.6 ± 0.64 |
| Reticulocyte Abs (/uL) | 466587.63 ± 35218.53 | 324936.55 ± 60280.44 | 367135.6 ± 59476.71 |
| Platelet Estimate (%) | Adequate | Adequate | Adequate |
| Neutrophils % | 84.25 ± 1.02 | 85.25 ± 1.77 | 82.6 ± 1.75 |
| Lymphocytes % | 14.63 ± 0.97 | 13.75 ± 1.78 | 16 ± 1.76 |
| Monocytes % | 1 ± 0.19 | 0.75 ± 0.25 | 1.4 ± 0.24 |
| Eosinophils % | 0.13 ± 0 | 0.25 ± 0.25 | 0 ± 0 |
| Basophils % | 0 ± 0 | 0 ± 0 | 0 ± 0 |
| RBC Morphology | Normal | Normal | Normal |
| Neutrophil Absolute | 2058.5 ± 253.87 | 1222.25 ± 184.14 | 1821.2 ± 197.2 |
| Lymphocytes Absolute | 327.88 ± 67.23 | 210.38 ± 54.81 | 367.6 ± 86.76 |
| Monocytes Absolute | 26.88 ± 6.02 | 9.88 ± 3.26 | 29.4 ± 3.57 |
| Eosinophils Absolute | 2.25 ± 0 | 2.38 ± 2.38 | 0 ± 0 |
| Basophil Absolute | 0 ± 0 | 0 ± 0 | 0 ± 0 |
| AST U/L | 215.25 ± 30.28 | 189.25 ± 25.95 | 186.2 ± 32.59 |
| ALT U/L | 36.13 ± 12.91 | 34.75 ± 3.75 | 19.8 ± 8.85 |
| Alkaline Phosphatase | 40.13 ± 0.99 | 44.25 ± 3.3 | 29.8 ± 2.54 |
| GGT U/L | 12.75 ± 2.31 | 13.13 ± 1.86 | 11.6 ± 0.81 |
| Total Bilirubin mg/dL | 0.24 ± 0.02 | 0.16 ± 0.03 | 0.16 ± 0.02 |
| Direct Bilirubin mg/dL | 0.05 ± 0.02 | 0.05 ± 0.03 | 0.03 ± 0.03 |
| Indirect Bilirubin mg/dL | 0.18 ± 0.03 | 0.05 ± 0.03 | 0.15 ± 0.03 |
| BUN mg/dL | 30.57 ± 1 | 29.38 ± 0.68 | 25.8 ± 1.43 |
| Creatinine mg/dL | 0.24 ± 0.03 | 0.2 ± 0.01 | 0.2 ± 0.03 |
| Cholesterol mg/dL | 210.5 ± 18.95 | 189.25 ± 11.65 | 204.8 ± 7.93 |
| Triglycerides mg/dL | 188.88 ± 45.5 | 75.88 ± 8.04 | 186.8 ± 41.55 |
| HDL mg/dL | 172 ± 13.02 | 172.25 ± 9.78 | 189.2 ± 11.96 |
| LDL mg/dL | 21.13 ± 3.47 | 14.25 ± 1.9 | 12.2 ± 1.16 |
| Uric Acid mg/dL | 3.43 ± 0.21 | 3.09 ± 0.29 | 3 ± 0.23 |
| Sodium mmol/L | 154.86 ± 1.73 | 153.4 ± 1.4 | 154.2 ± 1.88 |
| Potassium mmol/L | 5.43 ± 0.25 | 4.9 ± 0.24 | 5.8 ± 0.38 |
| Chloride mmol/L | 107.57 ± 0.95 | 109.2 ± 0.86 | 110.6 ± 1.81 |
| Carbon<br>Dioxide mmol/L | 13.19 ± 0.71 | 13.1 ± 0.81 | 15.18 ± 1.36 |

