## Supplementary figures and images for "Niclosamide Prodrug Enhances Oral Bioavailability and Targets Vasorin-TGFβ Signaling in Hepatocellular Carcinoma"

### Supplemental figures

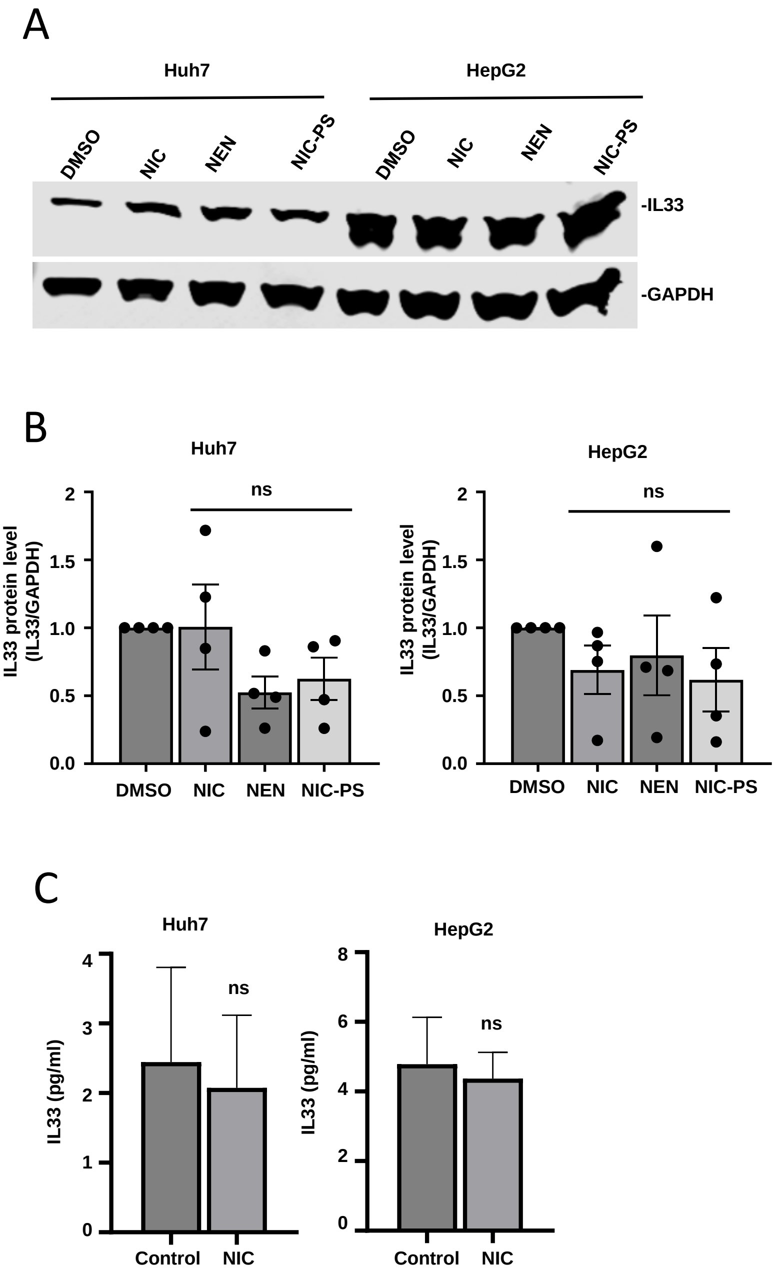

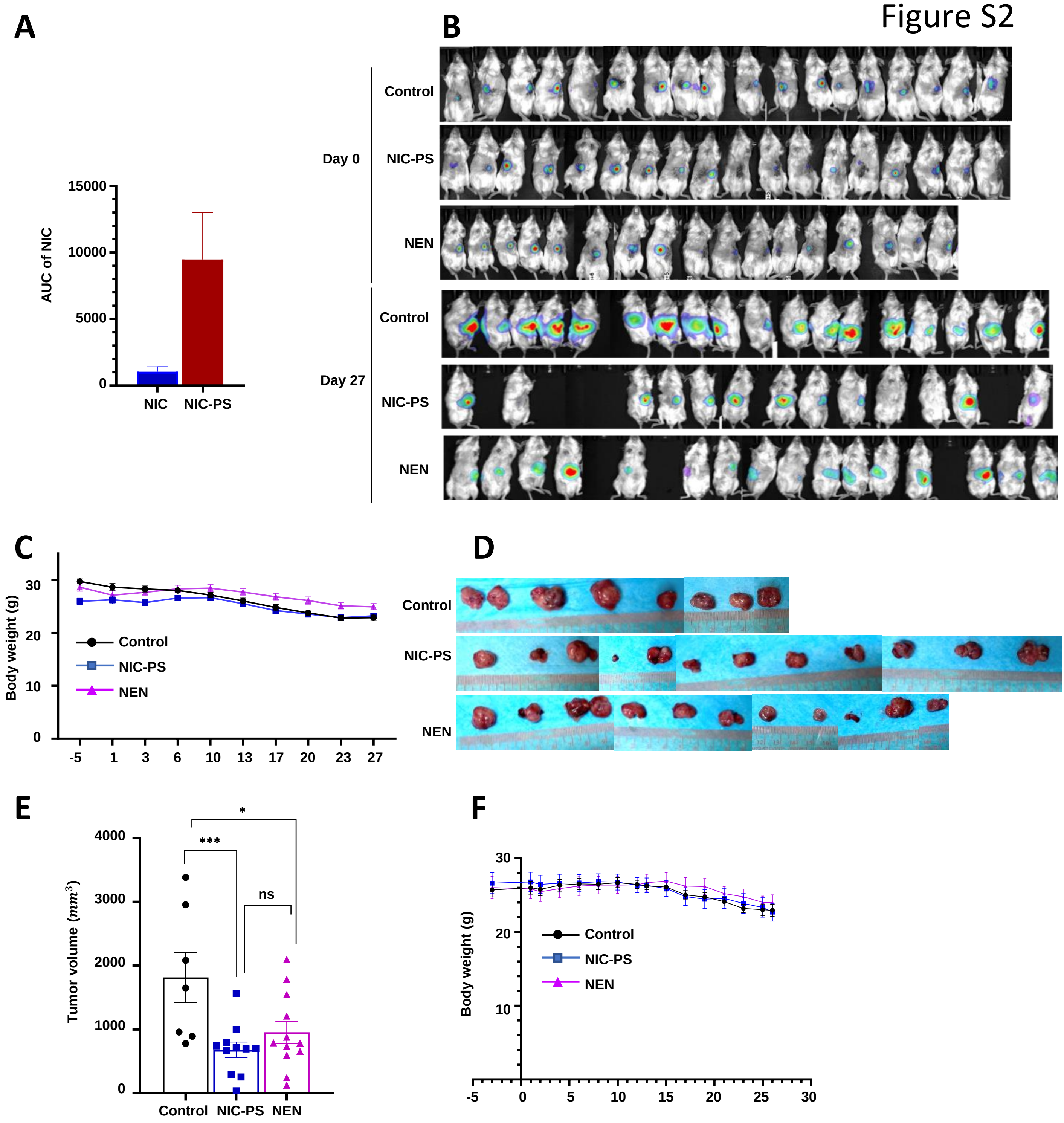

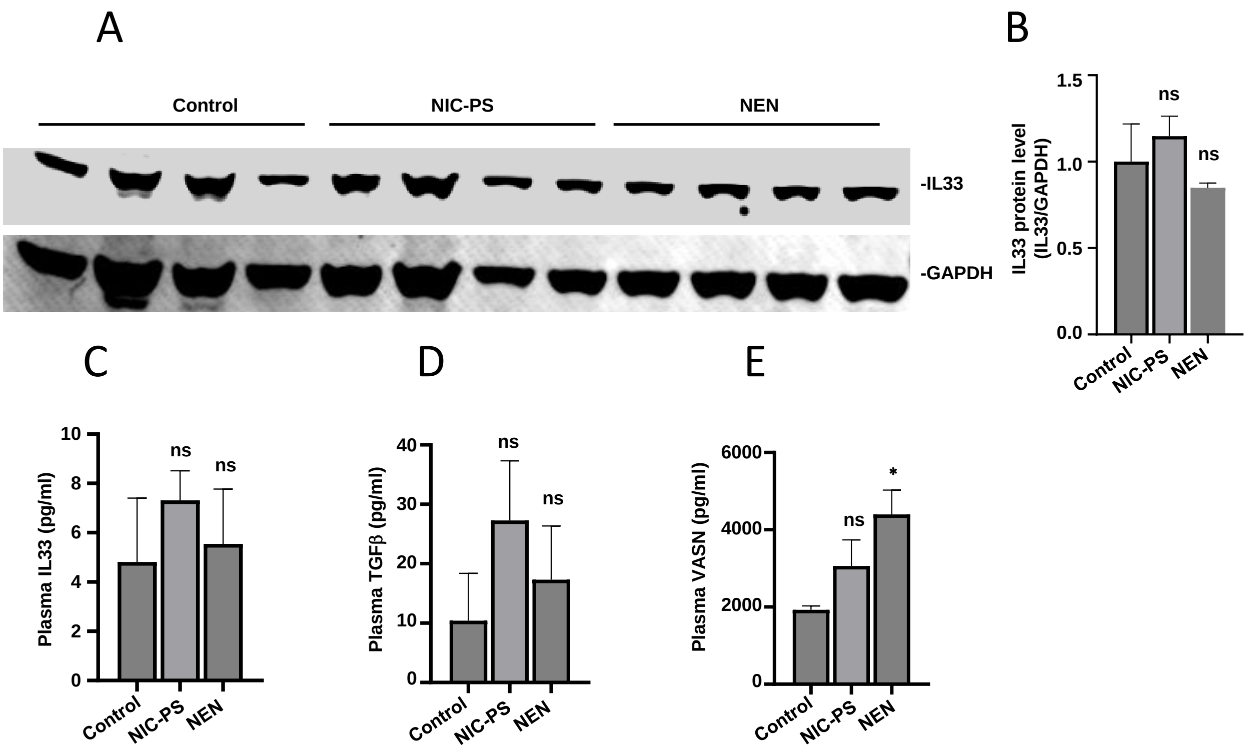

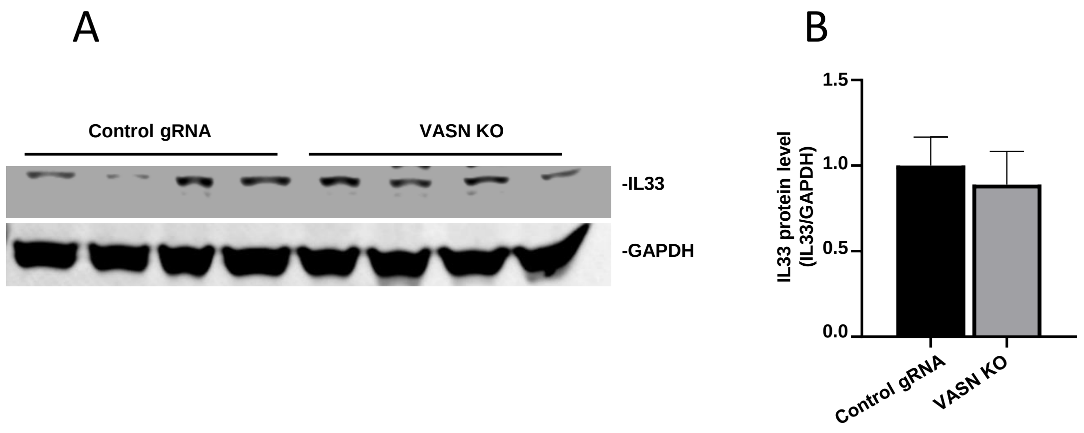
