## Supplemental materials for "Niclosamide Prodrug Enhances Oral Bioavailability and Targets Vasorin-TGFβ Signaling in Hepatocellular Carcinoma"

##### 1. Department of Surgery, School of Medicine, Asian Liver Center, Stanford, CA 94305, USA

##### 2. Current address: Guangzhou JOYO Pharmaceutical CO. LTD, Guangzhou 440112, China

3. Current address: Shanghai CureGene Pharmaceutical Co, Ltd., Shanghai 201203, China

\# Corresponding authors: Mingdian Tan, 1201 Welch Road, Palo Alto, CA 94305, United States, 1-650-724-3525,; Mei-Sze Chua, 780 Welch Road, Palo Alto, CA 94305, United States, 1-650-724-8601,

### **Supplementary Figure Legends**

**Figure S1. IL33 protein levels after treatment with NIC, NEN, or NIC-PS. A).** Western blot detection of IL33 protein level in Huh7 and HepG2 cells after treatment with NIC, NEN, or NIC-PS, and their **B).** quantified data. **C).** ELISA detection of IL33 protein in the cell supernatant, after treatment with NIC. Data are presented as Mean ± SEM, ** denotes significance at p-values < 0.01, in comparison to the control group.

**Figure S2. NIC-PS inhibited orthotopic HCC PDX xenograft growth.** **A)**. Blood AUC of NIC after oral administration of NIC or NIC-PS at dose equivalents to 40 mg/kg of NIC. **B)**. Monitoring of orthotopic PDX using bioluminescence imaging throughout the treatment period with NEN (200 mg/kg) or NIC-PS (100 mg/kg based on molecular weight equivalent to NIC). Compounds were administered daily by oral gavage. **C)**. Monitoring of mice body weight throughout the treatment period. In a second independent experiment, orthotopic PDX mice were treated with NEN and NIC-PS as in the first independent experiment. **D).** Harvested tumors and their tumor volumes **(E)** from the second experiment, after 4 weeks treatment with distilled water (untreated), NIC-PS, or NEN for 4 weeks; (n=8 to 12). **F)**. Monitoring of body weight changes in the second experiment. Data are presented as mean ± SEM; significance levels are denoted as *, and ** for p-values < 0.05, and <0.01, respectively, in comparison to the control group.

**Figure S3. Effect of NIC-PS and NEN on IL33, TGFβ, and VASN levels in treated mic.** **A).** IL33 protein level in tissue lysate and their **B).** quantified results. **C to E)**. Secreted IL33, TGFβ, and VASN protein levels in mice plasma collected at the last day of treatment, n=3 to 9. Data are presented as mean ± SEM.

**Figure S4.** **IL33 protein levels in Huh7 VASN KO xenografts.** **A).** Western blot detection of IL33 protein level in Huh7 control gRNA and Huh7 VASN KO xenografts, and their **B).** quantified data.
